# Relationships between the isotopic signatures of dissolved inorganic carbon and environmental variability in Northern Ireland coastal waters

**DOI:** 10.64898/2026.11.17.767286

**Authors:** William Ross Hunter, Claire Smyth

## Abstract

The isotopic signatures of dissolved inorganic carbon (δ^13^C-DIC) can provide a sensitive tracer of biological and biogeochemical processes in aquatic environments. However, isotopic measurements remain uncommon within routine coastal monitoring programmes. This pilot study investigated spatial variability in δ^13^C-DIC in surface waters across twelve Northern Ireland coastal monitoring stations sampled during July 2020. We tested the relationships between δ^13^C-DIC, dissolved inorganic carbon (DIC) concentration, nutrients, chlorophyll-a, temperature and salinity. δ^13^C-DIC exhibited substantial spatial variability, ranging from 1.25 to 2.53‰. Significant positive relationships were observed between δ^13^C-DIC and temperature (R^2^ = 0.55, p = 0.009), chlorophyll-a concentration (R^2^ = 0.38, p = 0.045) and DIC concentration (R^2^ = 0.52, p = 0.012), while significant negative relationships occurred with dissolved inorganic nitrogen (R^2^ = 0.53, p = 0.011) and phosphate (R^2^ = 0.44, p = 0.026). No relationships were detected between δ^13^C-DIC and salinity, suggesting that conservative mixing processes exerted little influence on isotopic variability across the study area. Bulk DIC concentration exhibited comparatively weak associations with environmental variables. The relationships we observed indicate that biological carbon fixation and nutrient use were important controls on spatial variation in δ^13^C-DIC in the coastal waters of Northern Ireland. These findings demonstrate the potential value of δ^13^C-DIC measurements as sensitive indicators of coastal carbon-cycle dynamics and support their inclusion within future coastal monitoring and assessment programmes.

**Research Highlights:**

- Here we present the first δ^13^C-DIC dataset reported for Northern Ireland coastal waters.
- Changes in δ^13^C-DIC was strongly correlated with differences in temperature, nutrients and chlorophyll-a across coastal sites.
- Isotopic enrichment of the DIC pool was consistent with biological carbon fixation and nutrient utilisation.
- Salinity explained little isotopic variability, indicating limited influence of conservative mixing processes.
- δ^13^C-DIC provided stronger environmental signals than bulk DIC concentration, highlighting its value for coastal carbon-cycle monitoring.

## 1. Introduction

Dissolved inorganic carbon (DIC) represents the largest readily exchangeable carbon reservoir in the ocean and plays an important role in regulating marine carbon cycling and air-sea CO_2_ exchange (Dai et al., 2022; Millero, 2007; Zeebe and Wolf-Gladrow, 2001). In seawater, DIC occurs predominantly as bicarbonate (HCO_3_^−^), with smaller contributions from carbonate (CO_3_^2−^) and dissolved CO_2_ species (Millero, 2007). Carbon occurs naturally as two stable isotopes, ^12^C and ^13^C, that differ in atomic mass but have the same chemical properties (Sulzman, 2007). The difference in atomic mass between ^12^C and ^13^C leads to subtle differences in reaction rates, causing isotopic fractionation during biological and geochemical processes such as photosynthesis and respiration (Zeebe and Wolf-Gladrow, 2001). As such, the isotopic composition of the marine DIC pool (δ^13^C-DIC) can be used to provide information on carbon sources and sinks, and the biogeochemical transformations occurring within an ecosystem (Bao et al., 2025; Cotovicz et al., 2019; Filipsson et al., 2017; Fry, 2006; Romanek et al., 1992; Zeebe and Wolf-Gladrow, 2001). Biological fixation of carbon during photosynthesis preferentially removes ^12^C leaving the residual DIC pool enriched in ^13^C. Consequently, changes in δ^13^C-DIC can provide insight into carbon cycling and primary production in marine ecosystems (Fry, 2006; Raymond and Bauer, 2001; Romanek et al., 1992). Despite its value as an integrative biogeochemical indicator, δ^13^C-DIC measurements remain comparatively scarce relative to routine observations of DIC, chlorophyll-a and inorganic nutrients. This lack of isotopic observations limits our ability to directly assess linkages between carbon fixation, ecosystem productivity and broader marine biogeochemical processes (Becker et al., 2012; Dai et al., 2022; Graven et al., 2021; Zeebe and Wolf-Gladrow, 2001).

Coastal waters are highly dynamic environments influenced by physical mixing, freshwater inputs, biological productivity and anthropogenic pressures, all of which can exert strong controls on dissolved inorganic carbon concentrations and isotopic composition (δ^13^C-DIC) (Fry, 2006; Graven et al., 2021; Zeebe and Wolf-Gladrow, 2001). Understanding the factors that drive variation in the coastal DIC pool is essential for resolving carbon cycling processes at the land-ocean interface, where exchanges between terrestrial, atmospheric and marine carbon reservoirs are particularly intense (Cotovicz et al., 2019; Dai et al., 2022; Raymond and Bauer, 2001). These processes are increasingly important in the context of climate change, biodiversity loss and chemical pollution, as coastal ecosystems are among the marine environments most exposed to multiple interacting stressors (Graven et al., 2021; Trégarot et al., 2024). Despite the potential value of δ^13^C-DIC measurements as indicators of ecosystem functioning and biogeochemical cycling (Becker et al., 2016; Becker et al., 2012; Graven et al., 2021; Zeebe and Wolf-Gladrow, 2001), they remain uncommon within routine marine monitoring programmes, which predominantly focus on nutrients, chlorophyll-a and hydrographic variables (Bauer et al., 2001; Becker et al., 2016; Sun et al., 2024). Developing the capacity to directly measure marine δ^13^C-DIC within routine monitoring programmes could improve our understanding of ecosystem conditions by enabling the detection of changes in coastal carbon cycling and biogeochemical functioning (Becker et al., 2016; Becker et al., 2012; Filipsson et al., 2017; Sun et al., 2024; Wang et al., 2026). Ultimately, this could strengthen evidence-based marine management and support regional assessments such as the OSPAR Quality Status Report (OSPAR, 2023a; b; c).

Northern Ireland is a constituent region of the United Kingdom with approximately 650 kms of coastline in the northeast of the island of Ireland. The water chemistry Northern Ireland’s coastal waters are routinely monitored by the Department of Agriculture, the Environment and Rural Affairs to meet the requirement of the UK Marine Strategy’s and EU Water Framework Directive’s programmes of measures for water quality, including indicators for eutrophication, chemical pollution and climate change. As part of these monitoring programmes the Agri-Food and Bioscience Institute Northern Ireland has conducted winter and summer coastal oceanography surveys since 2010, generating a high-quality observation of dissolved nutrients (ammonia, phosphate, oxidised nitrogen and silicate), chlorophyll and the underlying hydrographic conditions. However, sampling and measurement of DIC concentration and δ^13^C-DIC have not routinely been conducted in Northern Ireland’s coastal seas. As such, their relationships with environmental variability remain poorly characterised and their potential to provide information on coastal ecosystem functioning in Northern Ireland’s coastal seas remains unrealised. This paper presents a pilot study exploring spatial variability in dissolved inorganic carbon isotopic signatures (δ^13^C-DIC) across a subset of Northern Ireland coastal monitoring stations, sampled in July 2020. It was hypothesised that stations exhibiting evidence of enhanced phytoplankton production would display elevated δ^13^C-DIC values, reflecting preferential uptake of ^12^C during photosynthesis and enrichment of the residual DIC pool in ^13^C. Consequently, positive relationships were expected between δ^13^C-DIC and indicators of primary production and biological activity, including temperature and chlorophyll-a concentration, while negative relationships were anticipated with dissolved nutrient concentrations.

## 2. Materials and Methods

Water samples were collected from a subset of stations within the Northern Ireland coastal Water Framework Directive (WFD) monitoring programme conducted by the Agri-Food and Biosciences Institute (AFBI) in July 2020, during RV Corystes cruise CO2820 (Figure 1 A). At each station, vertical profiles of physical properties were obtained using a Sea-Bird 911 CTD equipped with a rosette water sampler. Near-surface water samples were collected in a 4 l Niskin bottle, using a Rosette water sampler. These water samples were recovered to the ship, filtered through sterile 0.22 μm membrane filters and transferred into Sterilin® containers. Samples were stored frozen at −20 °C until analysis.

**Figure 1.**
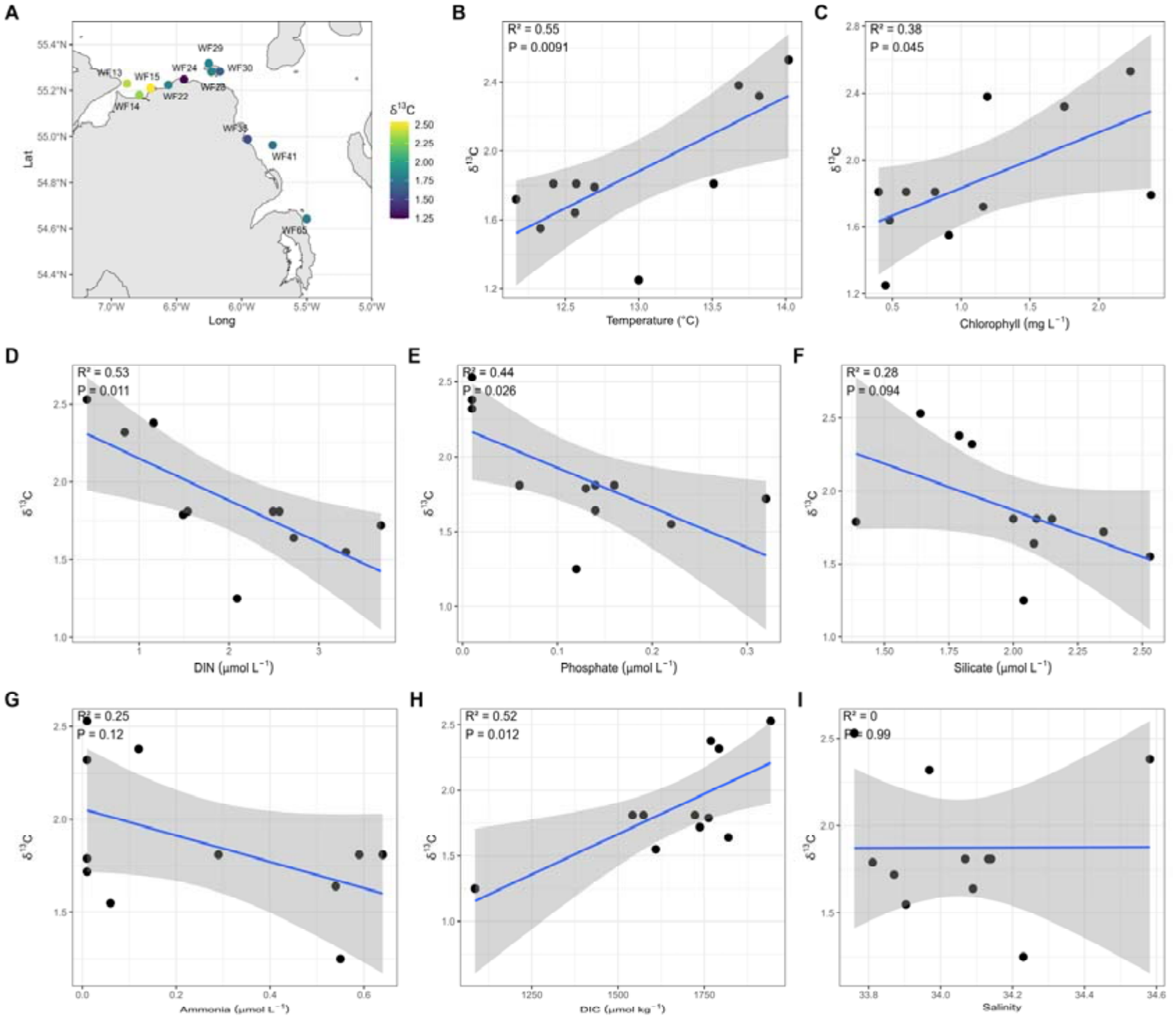
Spatial variability in dissolved inorganic carbon isotopic signatures (δ^13^C-DIC) across Northern Ireland coastal monitoring stations. (A) Sampling station locations coloured according to δ^13^C-DIC values (‰). Relationships between δ^13^C-DIC and environmental variables associated with biological production and nutrient availability, including (B) temperature, (C) chlorophyll concentration, (D) dissolved inorganic nitrogen (DIN), (E) phosphate, (F) silicate, (G) ammonia, (H) dissolved inorganic carbon concentration (DIC), and (I) salinity. Blue lines represent fitted linear regression models and grey shading denotes 95% confidence intervals. R^2^ and p-values are shown within panels.

Dissolved inorganic nutrients were determined at the AFBI Marine Chemistry Laboratory using a SEAL Analytical QuAAtro segmented-flow autoanalyzer following standard colorimetric methods. Ammonium was measured using the Berthelot reaction, while dissolved inorganic nitrogen (DIN) was determined as the sum of nitrate and nitrite following cadmium reduction and diazotisation. Dissolved phosphate and silicate were quantified using established colorimetric procedures. Nutrient concentrations are reported as μmol L^−1^ (Following Cheeseman and Wilson, 1978).

Samples for dissolved inorganic carbon (DIC) concentration and carbon isotope analysis (δ^13^C-DIC) were filtered through sterile 0.22 μm filters, transferred to Exetainer® vials (Labco, UK), and stored at 4 °C until analysis. Samples were acidified and converted to CO_2_ prior to determination of DIC concentration and isotopic composition using a GasBench II interface coupled to a DeltaPlus Advantage isotope ratio mass spectrometer (Thermo Finnigan, Bremen, Germany). Dissolved inorganic carbon concentrations were calculated from calibration curves generated using sodium carbonate standards, while isotopic compositions are reported in standard δ notation relative to Vienna Pee Dee Belemnite (VPDB).

Relationships between δ^13^C-DIC, DIC concentration, nutrients, chlorophyll a, temperature and salinity were examined using simple linear regression. Given the limited sample size (n = 12), multivariate analyses were not considered. To assess the influence of a potentially influential observation, selected analyses were repeated following exclusion of station WF24. All statistical analyses and figure production were undertaken in R (v4.4.3) using the tidyverse, sf, patchwork, and ggrepel packages (Pebesma, 2018; Pedersen, 2024; Slowikowski, 2024; Team, 2026; Wickham, 2016; Wickham et al., 2019).

## 3. Results and Discussion

δ^13^C-DIC exhibited clear spatial variability across the twelve Northern Ireland coastal stations sampled, ranging from approximately 1.2 to 2.6‰ (Figure 1A). δ^13^C-DIC increased significantly with seawater temperature (R^2^ = 0.55, p = 0.009; Figure 1B) and chlorophyll concentration (R^2^ = 0.38, p = 0.045; Figure 1C), with higher isotopic signatures observed at warmer and more productive stations. Significant negative relationships were observed between δ^13^C-DIC and dissolved inorganic nitrogen (DIN; R^2^ = 0.53, p = 0.011; Figure 1D) and phosphate concentrations (R^2^ = 0.44, p = 0.026; Figure 1E), while weaker negative trends were evident for silicate (R^2^ = 0.28, p = 0.094; Figure 1F) and ammonia (R^2^ = 0.25, p = 0.12; Figure 1G). A significant positive relationship was also observed between δ^13^C-DIC and dissolved inorganic carbon concentration (R^2^ = 0.52, p = 0.012; Figure 1H). No relationship was detected between δ^13^C-DIC and salinity (R^2^ < 0.001, p = 0.99; Figure 1I). In contrast to δ^13^C-DIC, dissolved inorganic carbon concentrations exhibited weak associations with environmental variables (Figure S1). Chlorophyll concentration explained the greatest proportion of observed DIC variability (R^2^ = 0.30, p = 0.082), whereas nutrient concentrations, temperature and salinity each explained less than 10% of the observed variance. Collectively, these relationships indicate that δ^13^C-DIC was most strongly associated with gradients in nutrient availability and conditions favourable for primary production across the study area. The positive association between δ^13^C-DIC and DIC concentration indicates that isotopic enrichment was not accompanied by a simple drawdown of the DIC pool. This may reflect the combined influence of biological production, carbon remineralisation and carbonate-system processes operating across the study area.

Our study provides the first assessment of spatial variability in δ^13^C-DIC across Northern Ireland coastal waters. The strongest relationships were observed with temperature, DIN and DIC concentration (R^2^ ≈ 0.5-0.6), with additional support from chlorophyll and phosphate. Collectively, these patterns indicate that δ^13^C-DIC was enriched at warm, chlorophyll-rich, nutrient-poor stations, consistent with biological carbon fixation influencing isotopic fractionation within the coastal DIC pool. Phytoplankton preferentially assimilate ^12^C during photosynthesis, enriching the residual DIC pool in ^13^C and increasing δ^13^C-DIC values. Similar relationships between δ^13^C-DIC, nutrient concentrations and indicators of primary production have been reported from the Skagerrak-Baltic region, where photosynthetic fractionation was identified as an important control on surface-water isotopic variability (Filipsson et al., 2017; Wang et al., 2026), and from productive freshwater reservoir systems (Bao et al., 2025).

Unlike indicators of biological activity and nutrient availability, salinity exhibited no meaningful relationship with δ^13^C-DIC. This suggests that freshwater-seawater mixing exerted relatively little influence on spatial isotopic variability during sampling. Such a result is perhaps unsurprising given the open and well-mixed nature of Northern Ireland’s surrounding shelf seas, in contrast to estuarine and river-influenced coastal environments where conservative mixing often represents a dominant control on δ^13^C-DIC variability (Raymond and Bauer, 2001). The positive relationship between δ^13^C-DIC and DIC concentration indicates that isotopic enrichment was not accompanied by a simple drawdown of the dissolved inorganic carbon pool. If biological uptake alone controlled spatial variability, enriched δ^13^C-DIC values would typically be expected to coincide with lower DIC concentrations as phytoplankton preferentially assimilate ^12^C during carbon fixation. However, DIC concentrations in coastal waters reflect the balance of multiple interacting processes, including biological production and respiration, air-sea CO_2_ exchange, carbonate-system reactions and physical transport. Consequently, isotopic enrichment of the DIC pool can occur without a proportional reduction in bulk DIC concentration (Filipsson et al., 2017; Ge et al., 2022). In these well-mixed coastal waters, biological uptake may therefore have modified the isotopic composition of dissolved inorganic carbon while physical transport and mixing replenished the DIC reservoir and offset local biological drawdown. Nevertheless, δ^13^C-DIC exhibited clear and coherent relationships with temperature, chlorophyll and nutrient concentrations, whereas DIC concentration itself showed comparatively weak environmental associations. These findings suggest that isotopic composition retained evidence of biological processing that was not readily apparent from concentration measurements alone, highlighting the potential value of δ^13^C-DIC as a sensitive tracer of coastal carbon-cycle dynamics. Similar conclusions have been drawn in both marine and freshwater systems, where δ^13^C-DIC has provided additional insight into carbon cycling processes beyond that available from bulk DIC measurements alone (Bao et al., 2025; Filipsson et al., 2017; Wang et al., 2026).

The small sample size in this pilot study means that the results should be interpreted with appropriate caution. We identified a potential outlier at station WF24, which exhibited notably low δ^13^C-DIC (1.25 ‰) and DIC concentration (1085.5 μmol kg^−1^) relative to the other twelve stations. Although chlorophyll-a concentration at WF24 was the second lowest recorded across the study area, the combination of depleted δ^13^C-DIC and unusually low DIC concentration was not consistent with the broader spatial patterns observed in the dataset. One possible explanation is analytical artefact associated with sample handling or storage (Deng et al., 2022). In particular, incomplete sealing of the Exetainer may have resulted in partial CO_2_ loss prior to isotope analysis. However, this explanation cannot be verified retrospectively. To assess the influence of WF24 on our observations, we reanalysed the data after excluding this station. Removal of WF24 strengthened two of the four principal relationships identified in the full analysis (Figure S2). The relationship between δ^13^C-DIC and temperature increased from R^2^ = 0.55 (p = 0.009) to R^2^ = 0.78 (p = 0.0006) (Figure S2A), while the negative relationship between δ^13^C-DIC and dissolved inorganic nitrogen (DIN) increased from R^2^ = 0.55 (p = 0.009) to R^2^ = 0.72 (p = 0.0019) (Figure S2B). Positive relationships between δ^13^C-DIC and DIC concentration (R^2^ = 0.36, p = 0.066) and chlorophyll-a (R^2^ = 0.31, p = 0.097) were also retained following exclusion of WF24, although explanatory power decreased for these variables (Figures S2C-D). The exclusion of WF24 did not alter the direction of any relationship observed in the dataset. The consistency in both direction and relative strength of the observed relationships suggests that the principal ecological interpretations are robust to the inclusion or exclusion of this potentially anomalous observation.

Despite the limited sample size, our study demonstrates spatial variability in δ^13^C-DIC across Northern Ireland coastal waters and identifies consistent relationships with temperature, nutrient availability and indicators of primary production. The stronger environmental relationships observed for δ^13^C-DIC compared with bulk DIC concentrations suggest that isotopic measurements may provide a useful insight into coastal carbon-cycle dynamics that cannot be captured by concentration data alone (Following Bao et al., 2025; Filipsson et al., 2017; Graven et al., 2021; Wang et al., 2026). The strong associations between δ^13^C-DIC, dissolved nutrient concentrations and chlorophyll-a indicate that carbon isotope signatures potentially serve as indicators of biological carbon fixation and primary producer activity in marine environments (Filipsson et al., 2017; Wang et al., 2026). However, δ^13^C-DIC measurements are not routinely collected within existing coastal monitoring programmes. Our findings suggest that stable carbon isotope observations could provide valuable complementary information for assessing coastal carbon cycling and ecosystem functioning. This interpretation is consistent with studies from both marine and freshwater systems, which have demonstrated the utility of δ^13^C-DIC as a tracer of biological and biogeochemical processes (Bao et al., 2025; Becker et al., 2016; Becker et al., 2012; Graven et al., 2021; Sun et al., 2024; Wang et al., 2026). Expansion of the present dataset across future years would allow the relative influence of biological production, nutrient cycling and physical mixing processes on coastal carbon dynamics to be investigated more comprehensively. These observations would strengthen our understanding of ecosystem processes and carbon cycling in the coastal waters of the Irish and Malin Seas and, thus, support future assessments of environmental status.

## 4. Conclusions

Spatial variability in δ^13^C-DIC was observed across twelve stations in Northern Ireland’s coastal waters, with isotopic signatures exhibiting consistent relationships with temperature, nutrient availability and indicators of primary production. The observed relationships with DIN, phosphate and chlorophyll-a suggest that isotopic variability was primarily linked to biological nutrient use and photosynthesis, and not to conservative mixing processes. Compared with bulk DIC concentration, δ^13^C-DIC exhibited stronger environmental relationships. Thus, demonstrating the potential value of δ^13^C-DIC measurements as an indicator of coastal carbon-cycle dynamics within future marine monitoring programmes.

## Acknowledgements

We acknowledge the assistance of Raymond Gilmore, Adam Mellor, Catherine Gibson, Carole Daly and Kirsty Whitehead in sample collection and processing. Stable Isotope analysis was carried out at the James Hutton Institute by Gillian Martin and Dr Barry Thornton. Sampling was conducted on the RV Corystes (Cruise No: CO2820) and we are grateful to the master and crew. This work was conducted under AFBI’s assigned work programme for the Department of Agriculture, the Environment and Rural Affairs (Northern Ireland).

## Data Availability

The data products, analytical workflows, and R scripts used in this study are archived in the Zenodo repository (DOI: 10.5281/zenodo.24310292) and are freely available from GitHub (<https://github.com/Billy-Hunter-AFBI/d13c-dic-northern-ireland-coastal-waters>). These resources enable full reproduction of the analyses and figures presented in the manuscript.

## Author Contributions

William R. Hunter: Conceptualization, Methodology, Formal analysis, Investigation, Data curation, Visualization, Writing - original draft, Writing - review & editing. Claire Smyth: Investigation, Resources, Data curation, Writing - review & editing.

## AI Use Statement

Microsoft Copilot was used to assist with language editing and the development of reusable analytical code for data processing, statistical analysis, and figure production. AI assistance was used primarily to improve code efficiency, automate repetitive workflows, and support the creation of publication-quality visualisations. All outputs were reviewed and validated by the authors, who take full responsibility for the accuracy, interpretation, and conclusions presented in this manuscript.

## Figures

**Supplementary Figure S1.**
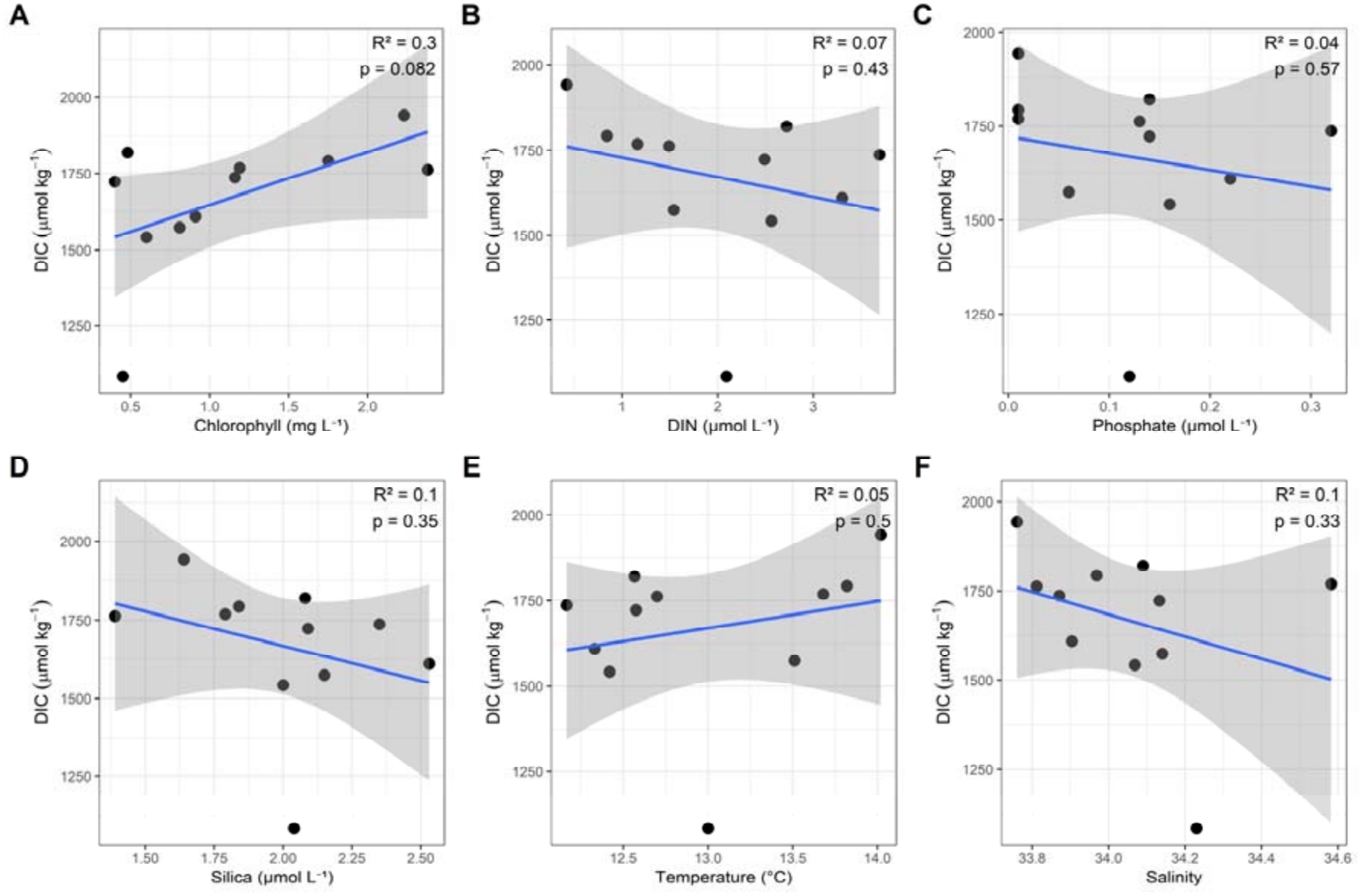
Relationships between dissolved inorganic carbon concentration (DIC; µmol kg^−1^) and environmental and biogeochemical variables in Northern Ireland coastal waters. **(A)** Chlorophyll concentration (mg L^−1^), **(B)** dissolved inorganic nitrogen (DIN; µmol L^−1^), **(C)** phosphate (µmol L^−1^), **(D)** silicate (µmol L^−1^), **(E)** temperature (°C), and **(F)** salinity. DIC exhibited a weak positive relationship with chlorophyll concentration, whereas relationships with nutrient concentrations, temperature and salinity were generally weak and explained relatively little of the observed variability. Solid blue lines represent fitted linear regressions and shaded regions denote 95% confidence intervals. Values of R^2^ and associated p-values are shown within each panel.

**Supplementary Figure S2.**
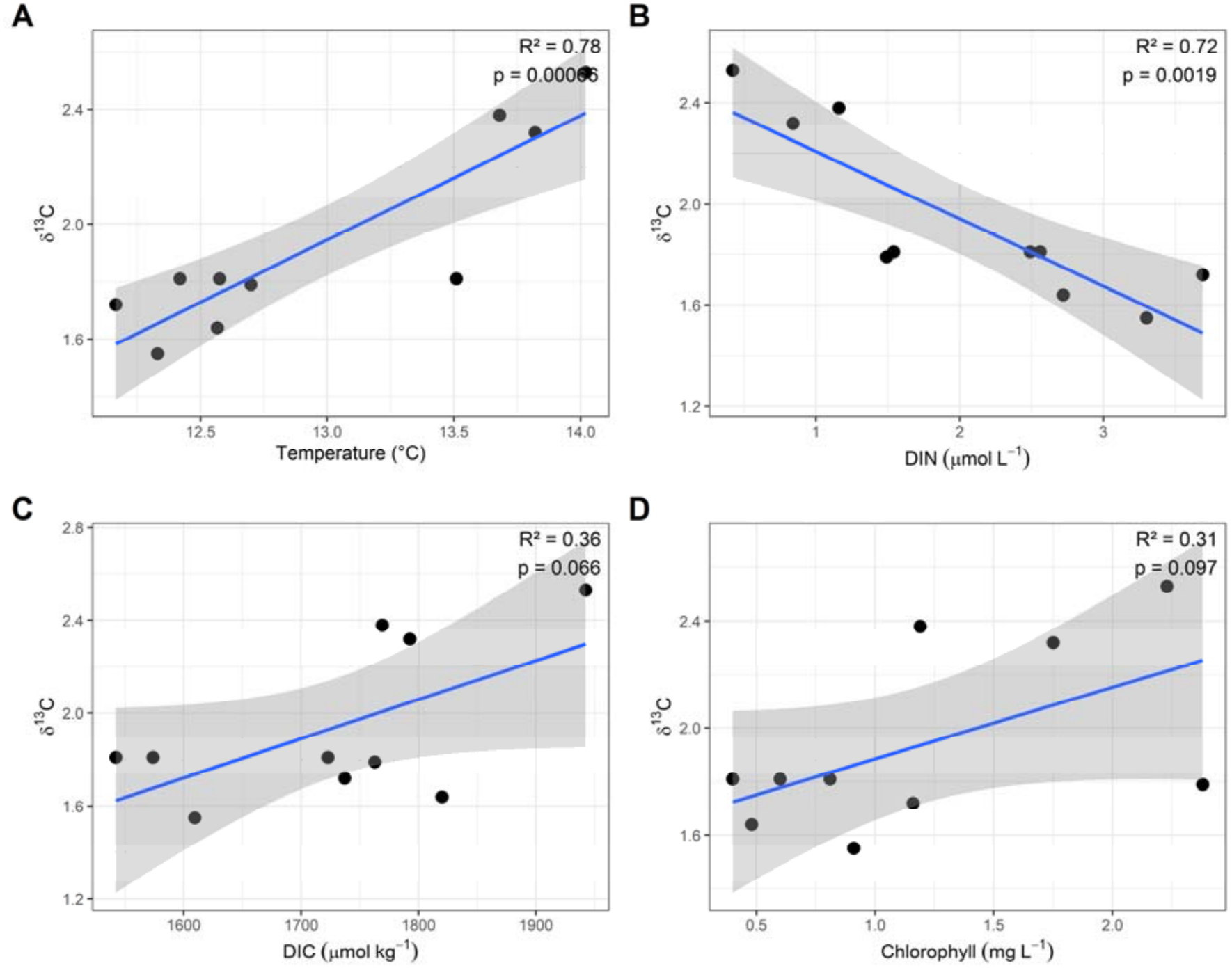
Relationships between dissolved inorganic carbon isotopic signatures (δ^13^C-DIC) and selected environmental and biogeochemical variables following exclusion of station WF24. **(A)** Temperature (°C), **(B)** dissolved inorganic nitrogen (DIN; µmol L^−1^), **(C)** dissolved inorganic carbon concentration (DIC; µmol kg^−1^), and **(D)** chlorophyll concentration (mg L^−1^). Exclusion of WF24 strengthened relationships between δ^13^C-DIC and both temperature and DIN, while positive relationships with DIC concentration and chlorophyll were retained. The consistency in the direction of these relationships indicates that the principal patterns observed in the full dataset were not dependent upon a single sampling station. Solid blue lines represent fitted linear regressions and shaded regions denote 95% confidence intervals. Values of R^2^ and associated p-values are shown within each panel.

